# Transcriptional and Spatial Resolution of Cell Types in the Mammalian Habenula

**DOI:** 10.1101/772376

**Authors:** Yoshiko Hashikawa, Koichi Hashikawa, Marcus L. Basiri, Yuejia Liu, Nathan L. Johnston, Omar R. Ahmad, Garret D. Stuber

**Affiliations:** Center for the Neurobiology of Addiction, Pain, and Emotion, Department of Anesthesiology and Pain Medicine, Department of Pharmacology, University of Washington, Seattle, WA 98195; Neuroscience Curriculum, University of North Carolina, Chapel Hill, NC 27599

## Abstract

The habenula complex is appreciated as a critical regulator of motivated and pathological behavioral states via its output to midbrain nuclei. Despite this, transcriptional definition of cell populations that comprise both the medial (MHb) and lateral habenular (LHb) subregions in mammals remain undefined. To resolve this, we performed single-cell transcriptional profiling and highly multiplexed *in situ* hybridization experiments of the mouse habenula complex in naïve mice and those exposed to an acute aversive stimulus. Transcriptionally distinct neuronal cell types identified within the MHb and LHb, were spatially defined, and differentially engaged by aversive stimuli. Cell types identified in mice, also displayed a high degree of transcriptional similarity to those previously described in zebrafish, highlighting the well conserved nature of habenular cell types across the phylum. These data identify key molecular targets within habenula cell types, and provide a critical resource for future studies.

## Introduction

The habenula complex serves as a neuroanatomical hub to regulate brain circuitry critical for many behavioral states related to motivation and decision-making (Hikosaka, 2010; Lecca et al., 2014; Namboodiri et al., 2016; Proulx et al., 2014; Viswanath et al., 2014). As part of the epithalamus, the habenula is well conserved across vertebrate species (Agetsuma et al., 2010; Amo et al., 2010; Bianco Isaac H and Wilson Stephen W, 2009; Lee et al., 2010; Stephenson-Jones et al., 2012), and in mammals, can be subdivided into medial (MHb) and lateral (LHb) subdivisions demarcated both anatomically and by patterned gene expression (Aizawa et al., 2012; Marburg, 1944). LHb neurons are almost entirely composed of *SLC17A6* (*VGlut2*) expressing glutamatergic neurons, which receive synaptic inputs from regions such as the ventral pallidum (Golden et al., 2016; Knowland et al., 2017), lateral hypothalamic area (Lazaridis et al., 2019a; Lecca et al., 2017; Stamatakis et al., 2016), endopeduncular nucleus/globus pallidus (Hong and Hikosaka, 2008; Li et al., 2019; Shabel et al., 2012; Wallace et al., 2017), the ventral tegmental area (VTA) (Root et al., 2014; Stamatakis et al., 2013), and other regions (Hikosaka et al., 2008). Glutamatergic LHb output neurons project to the VTA where they directly and indirectly regulate the activity of dopaminergic neurons to alter appetitive and aversive behaviors (Hong et al., 2011; Jhou et al., 2009; Lammel et al., 2012; Stamatakis and Stuber, 2012). Additionally, LHb neurons also project to the dorsal raphe to regulate the serotonergic system and depression-related behavioral phenotypes (Amo et al., 2014; Andalman et al., 2019; Li et al., 2011, 2013; Neckers et al., 1979; Wang and Aghajanian, 1977; Zhou et al., 2017).

Despite its close proximity, the MHb differs dramatically from the LHb with respect to its circuit connectivity, cellular composition, and function. The MHb contains spatially defined glutamatergic and cholinergic neurons (Aizawa et al., 2012), which receive synaptic input from the septum (Viswanath et al., 2014), and largely project to the interpeduncular nucleus (Herkenham and Nauta, 1979), where acetylcholine, glutamate, and other transmitters are released (Claudio Cuello et al., 1978; Grady et al., 2009; Molas et al., 2017; Ren et al., 2011a). While less is known about the function of MHb circuitry in controlling behavior, MHb neurons play a critical role in regulating nicotine reinforcement (Fowler and Kenny, 2012; Fowler et al., 2011; Lester and Dani, 1995), novelty preference (Molas et al., 2017), and anxiety/fear related behaviors (Hsu et al., 2014, 2016; Yamaguchi et al., 2013; Zhang et al., 2016). Collectively, the MHb and LHb represent two largely independent subsystems both of which importantly contribute towards regulating a variety of motivated behavioral states.

Despite the critical role of the habenula complex in orchestrating behavior, a comprehensive understanding of the genetically defined cell types in mammals within this region is still lacking. We used single cell RNA sequencing (Macosko et al., 2015) and multiplex *in situ* hybridization to generate a comprehensive dataset on the gene expression profiles of identified MHb and LHb cell types. Using unbiased clustering (Stuart et al., 2019) of transcriptomes from 11,878 cells, we identify 20 major cell types within the habenula complex, which can be further segregated into 12 neuronal subclusters from 5,558 cells. MHb and LHb neurons display distinct gene expression profiles for transcription factors, ion channel subunits, neurotransmitter systems, and G-protein coupled receptors, which provides novel genetic entry points for cell type specific targeting, as well as a resource of druggable targets that could act in cell type specific fashion. Single-cell transcriptional profiling of the habenula in zebrafish recently revealed a multitude of unique spatially and transcriptionally defined cell types (Pandey et al., 2018), which may share common features with mammals. By integrating our data with the previously published larval zebrafish dataset (Pandey et al., 2018), we show that habenula cell types from fish and mice show a striking degree of transcriptional similarity despite unique brain connectivity patterns between the species. Using highly-multiplexed *in situ* hybridization, we created a high-resolution atlas of habenular cell type distribution in the MHb and LHb. Finally, we identify at least two habenula cell types that are selectively activated, based on immediate early gene (IEG) expression, following exposure to an acute aversive stimulus, which highlights the functional diversity across genetically defined habenula cell types. Collectively, these data provide a comprehensive resource for future studies investigating habenular circuit and cellular function.

## Results

### scRNAseq identifies transcriptionally distinct clusters among neuronal and non-neuronal cells

To characterize transcriptional identities of single cells in the mammalian habenula, we utilized droplet (Macosko et al., 2015; Rossi et al., 2019) based chromium technology developed by 10x Genomics (Zheng et al., 2017). We microdissected the habenula from brain live slices of adult male mice (P50-55), enzymatically digested the tissue to obtain well dissociated highly viable single cells (viability >80 %), captured single cell transcriptomes, and generated cDNA libraries for subsequent sequencing (Figure 1). After computationally removing doublets (4.0 %) (DePasquale et al., 2018), we recovered the transcriptomes from a total of 11,878 cells with 17,726 genes in total (median unique molecular identifiers (UMIs)/cell: 2,030, median genes/cell: 1,001, Figure S1). In order to minimize the effects of experimental batch effects and behaviorally induced transcripts (e.g. IEGs), a computational technique (Stuart et al., 2019) to integrate multiple datasets by implementing both canonical correlation analysis (CCA) (Butler et al., 2018) and mutual nearest neighbor analysis (MNN) (Haghverdi et al., 2018) was utilized. Dimensions of the integrated data were reduced by principle component analysis followed by graph-based clustering (Macosko et al., 2015; Stuart et al., 2019) and visualization using the UMAP algorithm (Becht et al., 2019; McInnes et al., 2018) (Figure 1B). This method identified 20 cell clusters which were then assigned to neuronal and non-neuronal cell types by their canonical gene markers (Macosko et al., 2015; Saunders et al., 2018; Wu et al., 2017; Zeisel et al., 2018) (Figure 1B and Figure 1C). *Stmn2* and *Thy1* were selectively expressed in neuronal clusters, which were detected in ∼ 50 % of all the cells (Figure 1C and Figure S1), *Opalin* and *Mog* marked oligodendrocyte clusters, *Slc4a4* and *Ntsr2* were highly expressed in astrocyte clusters, and *Pdgfra*, *Gpr17* or *Ccnd1* were enriched in the oligodendrocyte precursor cells. This clustering method also detected less abundant cell types. *Tmem119* and *C1qc* clearly marked microglia (2.6 % of the total cells), *Tagln* marked the mural cell cluster (2.1 %), *Cldn5* and *Flt1* marked the endothelial cells (1.4 %) and *Fam216b* was exclusively expressed in the ependymal cluster (0.35 %).

**Figure 1.**
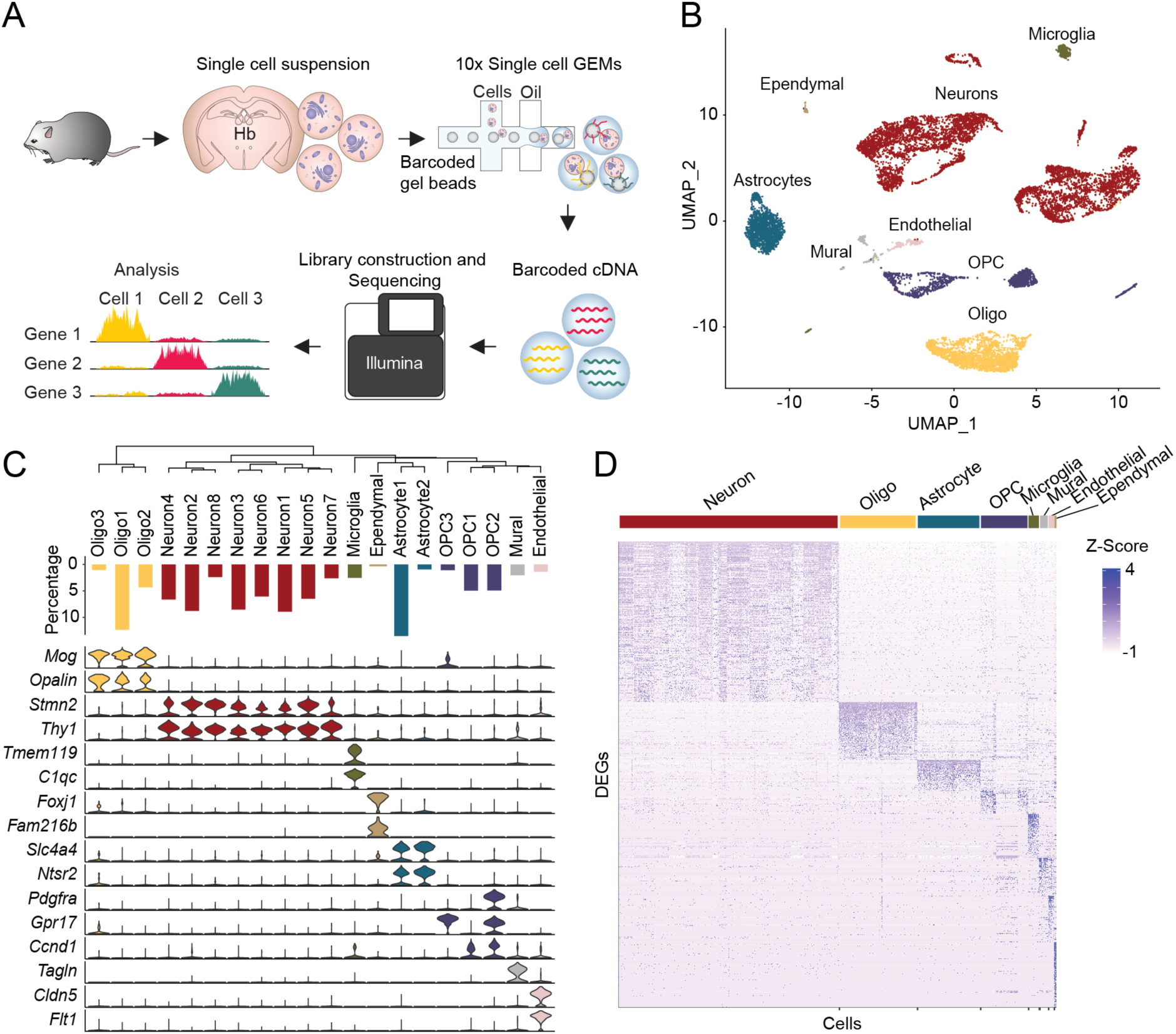
Single cell RNAseq analysis on mammalian habenula. **A.** Schematic of scRNAseq experiments. **B.** UMAP dimensional reduction and visualization of transcriptional profiles of 11,878 habenular cells. **C.** Top: dendrogram showing relationships between clusters. Middle: proportions of cells in each cluster. Bottom: violin plots showing levels of expression of canonical marker genes in neuronal and non-neuronal cells. **D.** Heat map showing scaled expression of all marker genes in each cell class (e.g. Neuron). Related to Figure S1.

Next, we calculated differentially expressed genes (DEGs) in the major cell types (e.g. Neuron, Astrocyte). A total of 1,449 genes were differentially expressed across the major cell types (corrected with a 5% false discovery rate (FDR) and log fold change > 0.25; Neuron: 405, Oligodendrocyte: 151, Astrocyte: 91, OPC: 82, Microglia: 138, Mural: 130, Endothelial: 230, Ependymal: 222). Interestingly, some detected DEGs were known habenula neuronal markers (e.g. *Tac2*, *Htr2c*) (Aizawa et al., 2012; Namboodiri et al., 2016) and markers of non-neuronal cells (*Kcnj10*, enriched in habenular astrocytes; *Cd68*, enriched in microglia) (Cui et al., 2018) (Valentinova et al., 2019) (Schmitt et al., 2012) (Figure 1D). To more broadly characterize the DEG expression profiles we assigned each DEG to specific functional domains for each cell type using gene ontology (GO) analysis (Chen et al., 2013; Kuleshov et al., 2016) (Figure S1). In Neuronal cells, the enriched GO terms pointed to processes critical for neural function in general (e.g. syntaxin-1 binding) (Zeisel et al., 2018) as well as terms indicative of habenular neurons (e.g. acetylcholine-gated cation-selective channel activity) (Ren et al., 2011). Collectively, these data broadly outline the cellular composition of the habenula complex and point to distinct cellular processes utilized by each cell type.

### scRNAseq identifies transcriptionally distinct clusters among habenular neurons

To further characterize neuronal cell types in detail, we re-clustered neuronal cells identified from the initial clustering. From this, we identified 16 discrete neuronal clusters (Figure 2A,B), among which 12 clusters were considered to be in the habenula due expression of canonical gene markers (Aizawa et al., 2012; Namboodiri et al., 2016) (see methods). Visualization of canonical gene markers in UMAP space readily separated habenular neurons into 6 MHb clusters (enriched with *Tac2*; 2,986 cells; median UMIs/cell = 3,750, median genes/cell = 1,840) and 6 LHb clusters (enriched in *Pcdh10*; 2,572 cells; median UMIs/cell = 3,483, median genes/cell = 1,822) (Figure 2B, C). Consistent with previous literature (Aizawa et al., 2012; Hsu et al., 2016; Namboodiri et al., 2016), the DEG analysis between MHb and LHb clusters revealed MHb specific (170 genes; *Slc17a7, Slc7a7, Tac2*) and LHb specific genes (289 genes; *Pcdh10*, *Htr2c*, *Gabra1*).

**Figure 2.**
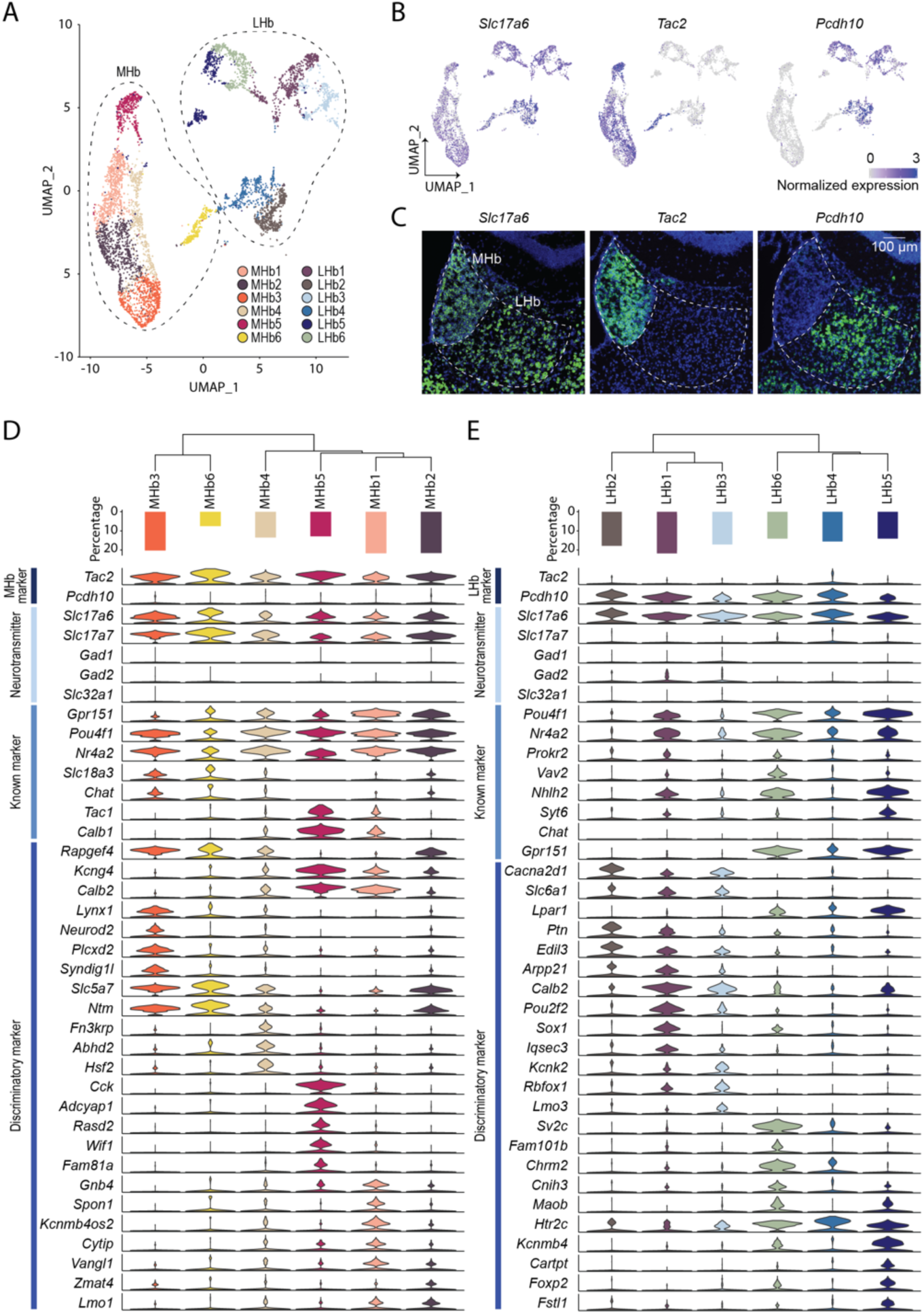
Transcriptional clusters of habenula neurons. **A.** UMAP visualization of 12 MHb or LHb neuronal clusters. 4 peri-habenula clusters were removed. **B.** Expression plots in UMAP space illustrating normalized expression values of *Slc17a6*, *Tac2* and *Pcdh10*. **C.** Representative FISH images for *Slc17a6*, *Tac2* and *Pcdh10* in and around the habenula. Scale bar: 100 µm. **D and E.** D and E correspond to the MHb and LHb respectively. Top: dendrogram showing relationships between clusters. Middle: proportions of cells in each cluster. Bottom: violin plots showing levels of expression of genes including entire MHb and/or LHb markers, major fast neurotransmitters, known markers and discriminatory markers in each cluster. Related to Figure S2-1 and S2-2.

To examine the transcriptional similarities between subclusters in the MHb and LHb, phylogenic trees were constructed based on a distance matrix in gene expression space (Figure 2D,E). In the MHb, subcluster #3 (MHb3) and #6 (MHb6) were closer in distance and were marked by *Slc18a3* and *Rapgef4* while MHb1, 2, 4 and 5 were in close proximity and were enriched with *Kcng4* and *Calb2*. We further examined whether each MHb subcluster had unique marker genes. 16-224 genes were enriched in each MHb subcluster but few single genes were cluster specific (expressed in >30 % of cells in the enriched cluster; expressed in <10% of cells in the other subclusters). Thus, the majority of DEGs did not distinguish individual clusters alone, but their expression contributed towards a unique transcriptional profile, which when analyzed in aggregate accurately defined distinct cell types (expressed in >10% of cells in the other subclusters). Notable selective genes, *Lynx1*/*Neurod2, Fn3krp*, *Cck*/*Adcyap1*, and *Kcnmb4os2* were expressed in MHb3, MHb4, MHb5 and MHb1 respectively (Figure 2D). Similarly, in the LHb, LHb1,2 and 3 were grouped with the expression of *Cacna2d1* and *Slc6a1* while LHb4,5 and 6 were more closely related and were marked with *Gpr151*, *Lpar1* and *Htr2c*. Differential gene expression analysis revealed 77-147 enriched genes in each LHb subcluster. In contrast to the MHb, the majority of LHb subclusters could only be defined by the combination of genes except for LHb4, which was enriched with *Fam101b* and *Sv2c* (Figure 2E). Taken together, most habenula neuronal cell types can only be defined by multiple, but not single genes.

### Distinct transcriptional domains between the MHb and the LHb

To better understand enrichment in functional domains of MHb or LHb enriched genes, we performed GO analysis on DEGs between the aggregated MHb and LHb neuronal data (the numbers of DEGs: 170 in the MHb; 289 in the LHb) (Chen et al., 2013; Kuleshov et al., 2016) (Figure 3). In the MHb, DEGs were particularly enriched in the cellular functions related to cholinergic transmission and membrane conductance (e.g. calcium ion binding, ion channels). Consistent with studies showing that the MHb is a major source of acetylcholine in the midbrain Interpeduncular nucleus (IPN) (Flumerfelt and Contestabile, 1982; Ren et al., 2011; Zhang et al., 2016), genes related to the machinery for cholinergic synthesis (*Chat*) and Ach gated ion channels (*Chrna3*, *Chrna4*, *Chrnb3*, *Chrnb4*) were also abundantly expressed (Figure 3 and Figure S3). Interestingly, we found enrichments of genes related to several types of potassium channels (e.g. *Kcna2*, *Kcnma1*, *Kcnip1*) and calcium ion binding (e.g. *Cadps2*, *Pcp4*, *Tesc*, *Syt15*), some of which were previously well characterized in slice electrophysiology studies (Hsu et al., 2014; Quina et al., 2009) (Figure 3 and Figure S3).

**Figure 3.**
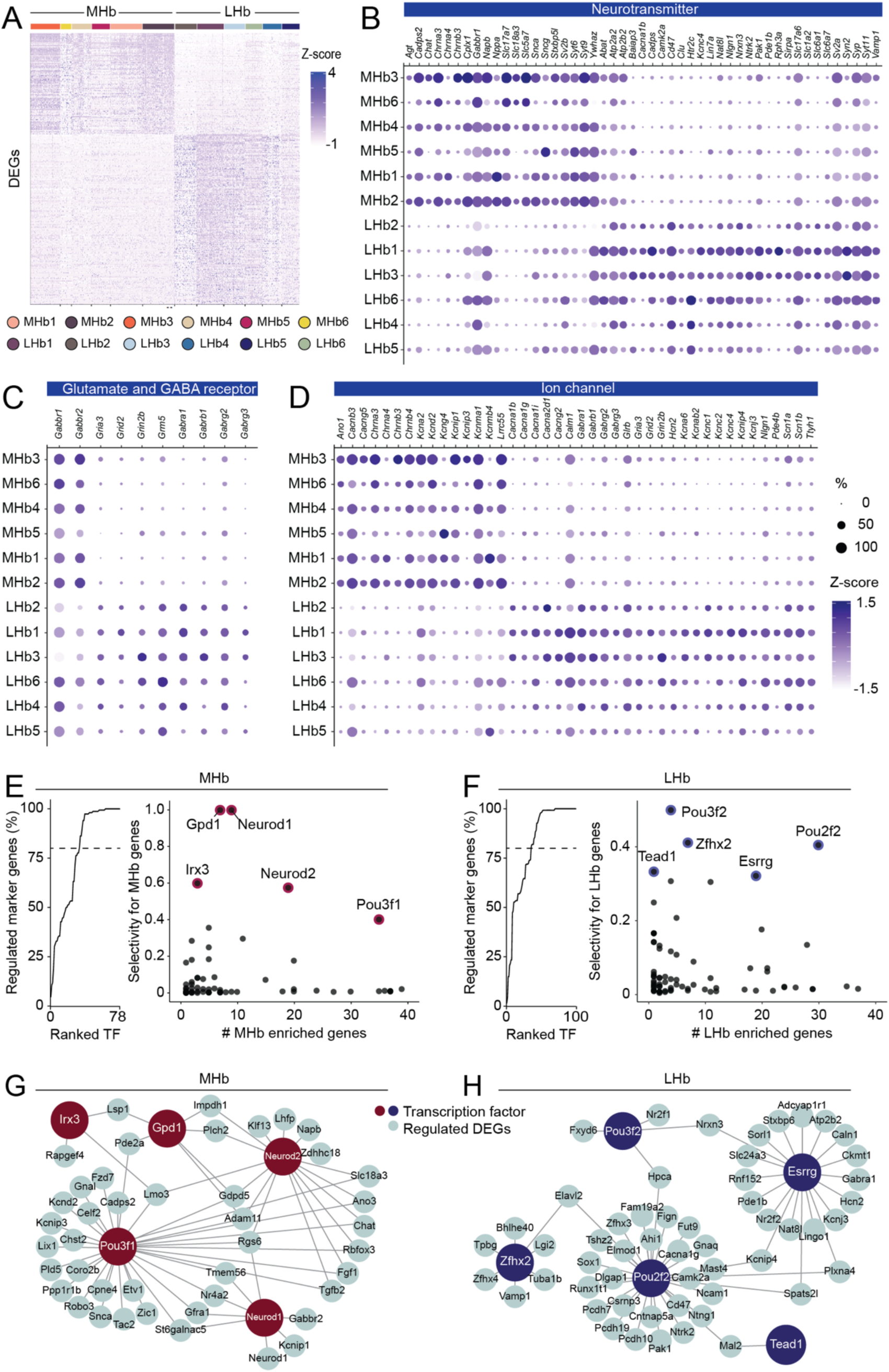
Comparative analysis of transcriptions and their regulations between the MHb and the LHb. **A.** Heat map showing scaled expression of all marker genes for entire MHb clusters or LHb clusters. **B, C and D.** Dot plot illustrating scaled expression levels (color) and the proportions of expressing cells (dot size) of Neurotransmitter related genes (**B**), Glutamate and GABA receptor related genes (**C**) and Ion Channel related genes (**D**) in each MHb and LHb cluster. **E and F.** Left: Cumulative percentage of MHb (**E**) or LHb (**F**) marker genes regulated by transcription factors (TF) sorted by their selectivity to regulate marker genes. Right: Scatter plot showing the number of marker genes regulated by transcription factors (only TFs regulating <40 markers are shown) and the selectivity of gene regulation for marker genes. 5 genes with highest selectivity scores were highlighted. **G and H.** Schematic illustrating the regulation of marker genes by 5 TFs of high selectivity in the MHb (**G**) and the LHb (**H**). Related to Figure S3.

Ontology analysis in the LHb pointed to functions in fast neural transmission (Glutamate and GABA) and membrane conductance as well. As the LHb is innervated heavily from the entopeduncular nucleus (EPN) co-releasing Glutamate and GABA (Herkenham and Nauta, 1979; Hong and Hikosaka, 2008; Lazaridis et al., 2019; Shabel et al., 2014) and the lateral hypothalamic area (LHA; both GABAergic and Glutamatergic innervations) (Lazaridis et al., 2019; Li et al., 2019), we found enrichments in genes that are components of GABA receptors (e.g. *Gabra1*, *Gabrb1*) and Glutamate receptors (e.g. *Gria3*, *Grid1*). In addition, we found enrichments in various types of voltage gated ion channels that critically define cellular electrophysiological properties including sodium channels (e.g. *Scn1a*, *Scn1b*), calcium channels (e.g. *Cacna1b*, *Cacna1g*) (Anderson et al., 2010), potassium channels (e.g. *Kcnc4*, *Kcna6*) and hyperpolarized-activated cyclic nucleotide-gated cation channels (*Hcn2*) (Poller et al., 2011) (Figure 3 and Figure S3).

### Distinct transcriptional controls between the MHb and the LHb

The divergent repertoires of gene expressions in the MHb and the LHb suggests that their expression is governed by distinct gene regulatory networks. To reconstruct gene regulatory networks in the MHb and the LHb from scRNAseq data, we performed SCENIC analysis, which utilizes both co-expression modules between transcription factors (TFs) and candidate target genes, as well as databases of DNA binding motifs of TFs to infer significant gene regulation by transcription factors (Aibar et al., 2017; Davie et al., 2018) (Figure 3E-H). SCENIC identified 78 and 100 TFs regulating MHb enriched and LHb enriched genes, respectively (Figure 3E,F). Since some TFs ubiquitously regulated gene transcription (e.g. Elf2 and Kdm5b, which regulate over 1,000 genes), we ranked the identified TFs by their selectivity to MHb or LHb enriched genes. We found that the top 33 TFs (out of 78) in the MHb and the top 36 (out of 100) in the LHb regulated over 80% of the enriched genes (Figure 3E, F) whereas only 9 of them (25%) were common between the two subregions. This suggests that enriched genes in the MHb and LHb are controlled by distinct gene regulatory networks. In addition, the MHb and LHb enriched genes were combinatorially regulated. The majority of enriched genes (82.4% in the MHb, 86.9% in the LHb) were regulated by multiple TFs (Figure 3G, H).

### Transcriptional conservation of habenular neurons between mice and fish

Previous studies suggest that homologous anatomical and molecular features in the habenula exist between mice and fish (Aizawa et al., 2005, 2012; Amo et al., 2010; Bianco and Wilson, 2009; Gamse et al., 2005; Herkenham and Nauta, 1979; Stephenson-Jones et al., 2012). To examine cross-species correspondence of the habenular transcriptome, we utilized Seurat V3 (Butler et al., 2018; Stuart et al., 2019) to jointly integrate the scRNAseq data set of adult zebra fish habenula neurons (Pandey et al., 2018) (performed with chromium technology; GSE105115) and our mouse neuron data (Figure 4A). Without integration, fish and mouse neuronal clusters were nearly non-overlapping (Figure 4B and Figure S4), Seurat integration and clustering identified 10 clusters (Integ1-10), each of which consisted of both mouse and fish neurons (Figure 4B,C and Figure S4). The expression of *Tac2*, a canonical marker for mouse MHb and zebrafish dorsal habenula (Aizawa et al., 2005; Gamse et al., 2005), was restricted in Integ2, 3, 5, 6 and 7 while *Pcdh10*, a marker for mouse LHb and zebrafish ventral habenula (Aizawa et al., 2012), marked remaining clusters (Figure 4D). To examine the extent of correspondence between integrated clusters with neural clusters in mice or zebrafish clusters (Figure S4), Pearson correlation coefficients were calculated (See methods). Consistent with canonical marker expressions, Integ2, 3, 5, 6 and 7 were moderately correlated with zebra fish dorsal habenula and mouse MHb clusters whereas Integ1, 4, 8, 9 and 10 corresponded to the ventral part or interneurons in zebrafish habenula and mouse LHb clusters (Figure 4E) with homologous gene expressions in corresponding clusters (Figure 4F), suggesting that in agreement with anatomical studies (Aizawa et al., 2005; Amo et al., 2010; Gamse et al., 2005; Herkenham and Nauta, 1979), the dorsal and ventral habenula of zebrafish are transcriptionally homologous to the MHb and the LHb of mice respectively.

**Figure 4.**
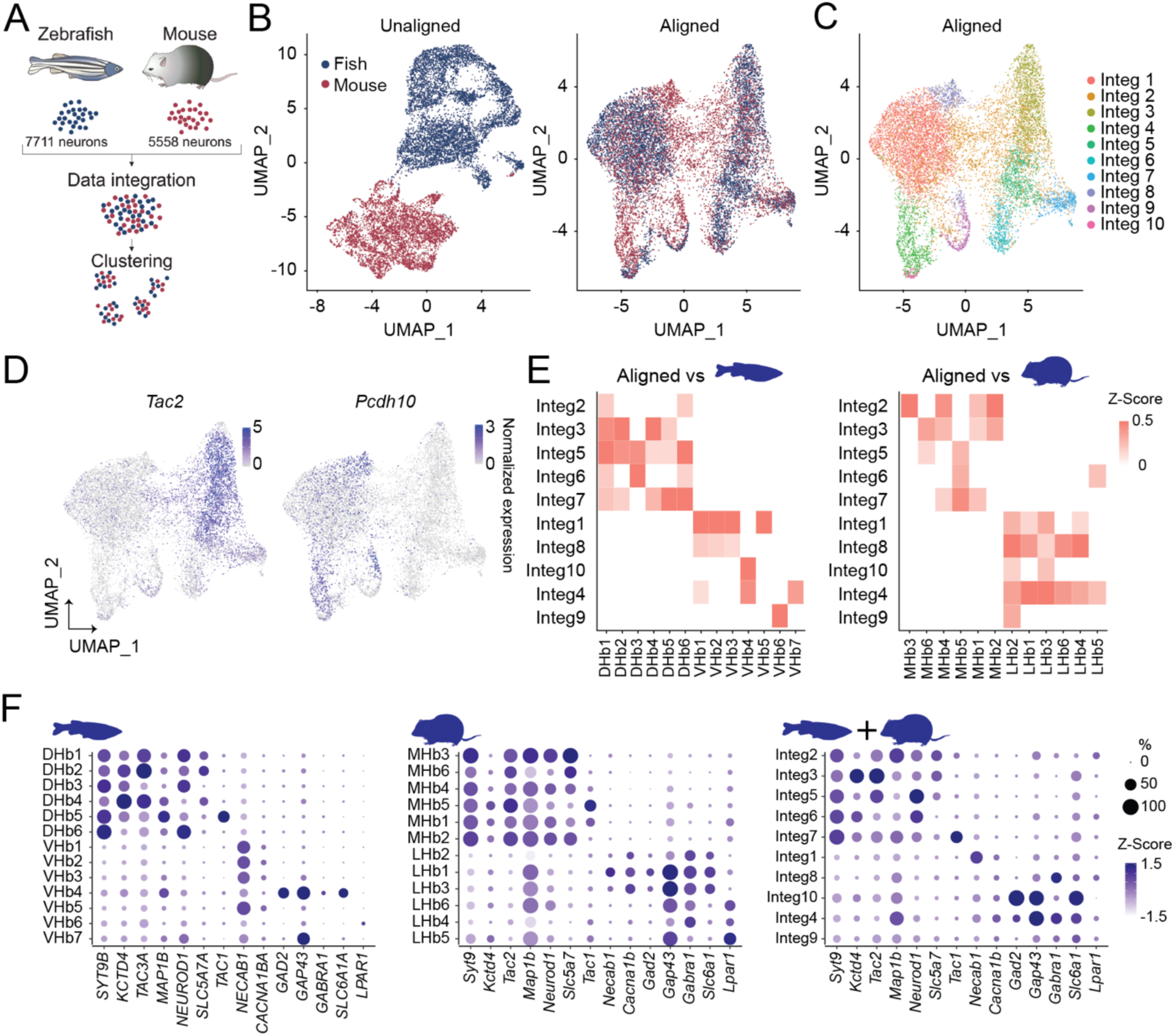
Integrative cross-species analysis on transcriptomes of mouse and fish habenula. **A.** Schematics illustrating integrative cross species analysis of mouse and zebrafish transcriptional profiles of habenula neurons. **B.** UMAP dimensional reduction of transcriptional profile of jointly analyzed zebrafish and mice habenula neurons without alignment (Left) and with Seurat V3 alignment (Right). **C.** UMAP visualization of 10 clusters of jointly analyzed neurons. **D.** Expression plots showing normalized expression values for *Tac2* and *Pcdh10* in the UMAP space. **E.** Heatmap illustrating Pearson correlations between integrated clusters and zebrafish clusters (Left) and between integrated clusters and mouse clusters (Right). White, p>0.05 or correlation value<0. **F.** Dot plot showing scaled expression levels (color) and the proportion of expressing cells (dot size) of marker genes in zebrafish (Left), mouse (Middle) and integrated (Right) clusters. Related to Figure S4.

### Visualization of transcriptionally defined cell types *in situ*

In order to cross-reference our scRNAseq data and visualize transcriptionally defined neuronal clusters in space, we performed Hiplex FISH developed by ACDbio, which permits detection of up to 15 genes within intact tissue (Figure 5A). 12 marker genes in the MHb or LHb were detected by sequential rounds of hybridization, amplification, imaging and cleavage (Figure 5A-C). Because we detected 6 MHb clusters in the scRNAseq data, we next used Seurat clustering on the 22,432 cells in the Hiplex dataset to partition the data into 6 clusters (hMHb1-6) (Figure 5B, D). Correlation analysis identified corresponding clusters between the scRNAseq and Hiplex datasets with moderate to high correlation coefficient values (0.48-0.95), with the exception that MHb4 did not have a corresponding Hiplex cluster presumably due to its lack of clear single gene markers. To examine the spatial distribution of the Hiplex clusters, anatomical coordinates of all cells were reconstructed from the raw microscopy images (Figure 5F and Figure S5). MHb clusters were latero-medially and dorso-ventrally biased and patterns of spatial distributions were consistent along the anterior-posterior axis. For instance, hMHb4, which was marked by *Neurod2*, *Zmat4* and *Synpr* and correlated with the scRNAseq cluster MHb3, was concentrated in the medio-ventral part of the MHb while hMHb5 and 6, which were enriched with *Kcng4* and *Cck* and corresponded with MHb5, were distributed in the dorsomedial and dorsolateral MHb (Figure 5D, F). Similar approaches were applied to the LHb. In the LHb, clustering based on the expression of 12 genes in 13,151 cells identified 6 clusters (hLHb1-6) (Figure 5C, E). Correlation analysis between all the cluster pairs of the two modalities identified corresponding clusters with weak to strong correlations (0.21-0.75). Similar to the MHb clusters, reconstruction of the spatial distribution of LHb Hiplex clusters revealed clear topographical organization (Figure 5G). In addition to dorso-ventral and latero-medial bias, hLHb clusters tended to show unique distributions on antero-posterior axis as well. hLHb2, which was enriched with *Necab1* and corresponded the most with LHb1 in the scRNAseq data, was most concentrated in the dorsal-medial part of the anterior LHb while hLHb3, which was marked by *Fam101b* and corresponded with LHb6, was biased in the ventral-lateral part of the middle LHb (Figure 5E, G). Thus, Hiplex FISH analysis along with correlation analysis with scRNAseq data revealed topographical organization of transcriptionally identified clusters in the habenula.

**Figure 5.**
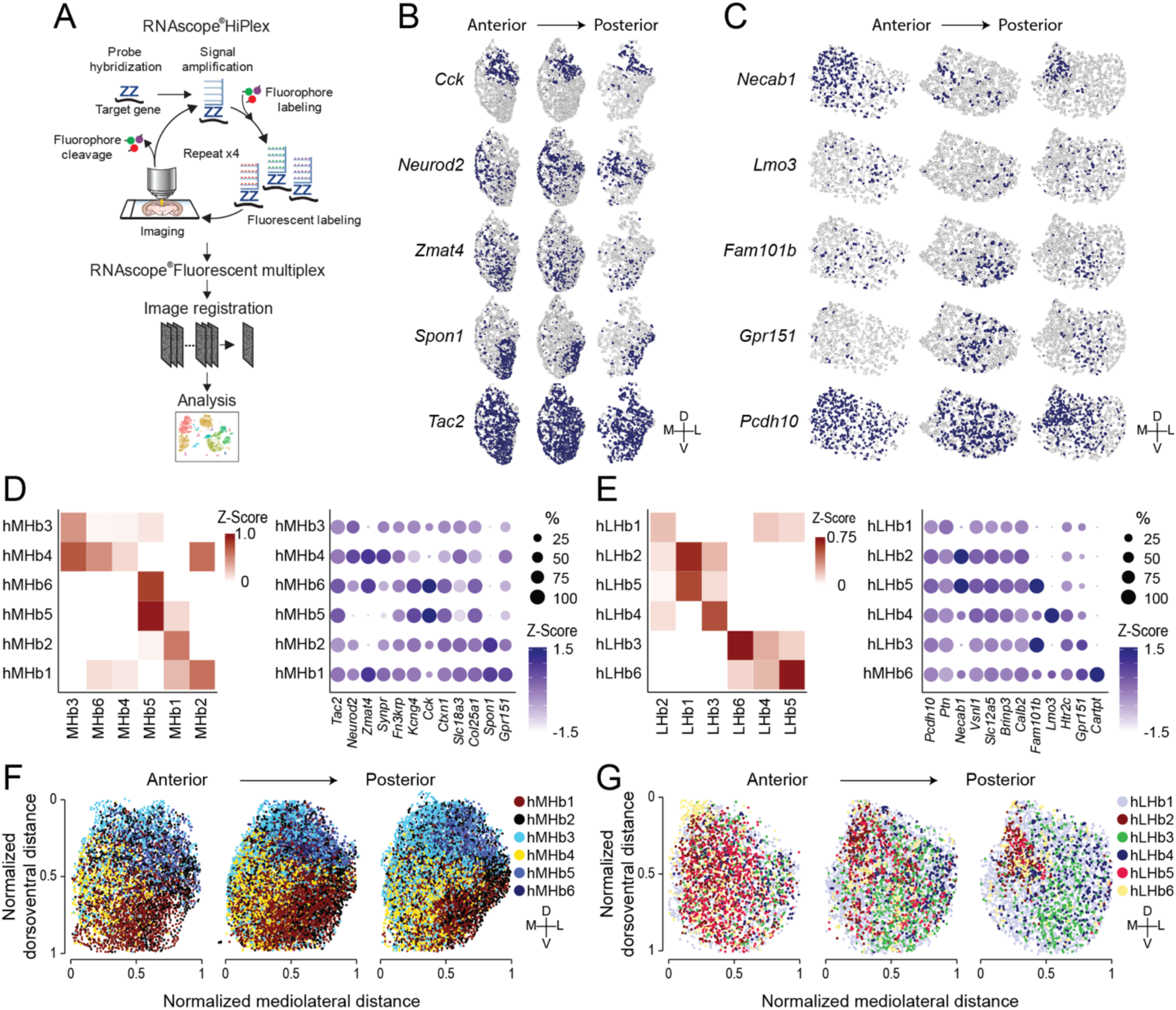
Visualization of habenular transcriptional cell types in situ. **A.** Schematic illustrating experimental design of Hiplex FISH. **B and C.** Representative ROI images showing *in situ* expressions of genes targeting the MHb (**B**) and the LHb (**C**). **D and E.** Left: Heatmap showing Pearson correlations between Hiplex clusters and scRNAseq clusters of MHb (**D**) and LHb (**E**). White, correlation value<0. Right: Dot plot illustrating the expression levels (color) and the proportions of expressing cells (dot size) of marker genes in MHb (**D**) and LHb Hiplex clusters (**E**). **F and G.** Spatial distributions of MHb (**F**) and LHb clusters (**G**) along anterior posterior axis.

### Identification of MHb and LHb subclusters responding to an aversive stimulus

Given that both the MHb and the LHb mediate a variety of behavioral responses to aversive and appetitive stimuli including stress, pain, and addictive compounds such as nicotine and cocaine (Lazaridis et al., 2019; Matsumoto et al., 2005; Stamatakis and Stuber, 2012; Velasquez et al., 2014; Zhang et al., 2016), we examined whether cellular activation to an aversive stimulus was transcriptionally detectable in the habenula, and if so, whether acute aversive stimulus exposure engaged transcriptionally defined cell types. Since we utilized Act-seq for our scRNAseq data acquisition (Wu et al., 2017), a technique for enhancing the signal to noise ratio of IEG detection, we could compare how acute foot-shock altered IEG expression. Experimental mice received 30 foot-shocks in a 60 min session while control mice remained in their home cages. We then compared IEG expression across all MHb and LHb neuronal subclusters, by examining the expression level and the proportion of *Fos* and *Egr1* between shock and control groups (Figure 6). This revealed that MHb2,4 and 5 showed a significant increase in *Fos* and/or *Egr1* expression while only MHb3 showed a significant increase in the percentage of *Fos* and *Egr1* (+) cells. For the LHb, LHb1, 3, and 6 showed increased *Fos* or *Egr1* expression, but only LHb6 also showed a significant increase in the proportion of *Fos* and *Egr1* (+) cells. Collectively, these results identify at least two transcriptionally defined and genetically accessible cell types (MHb3 and LHb6) that are robustly engaged following foot-shock exposure, which can be prioritized for future study.

**Figure 6.**
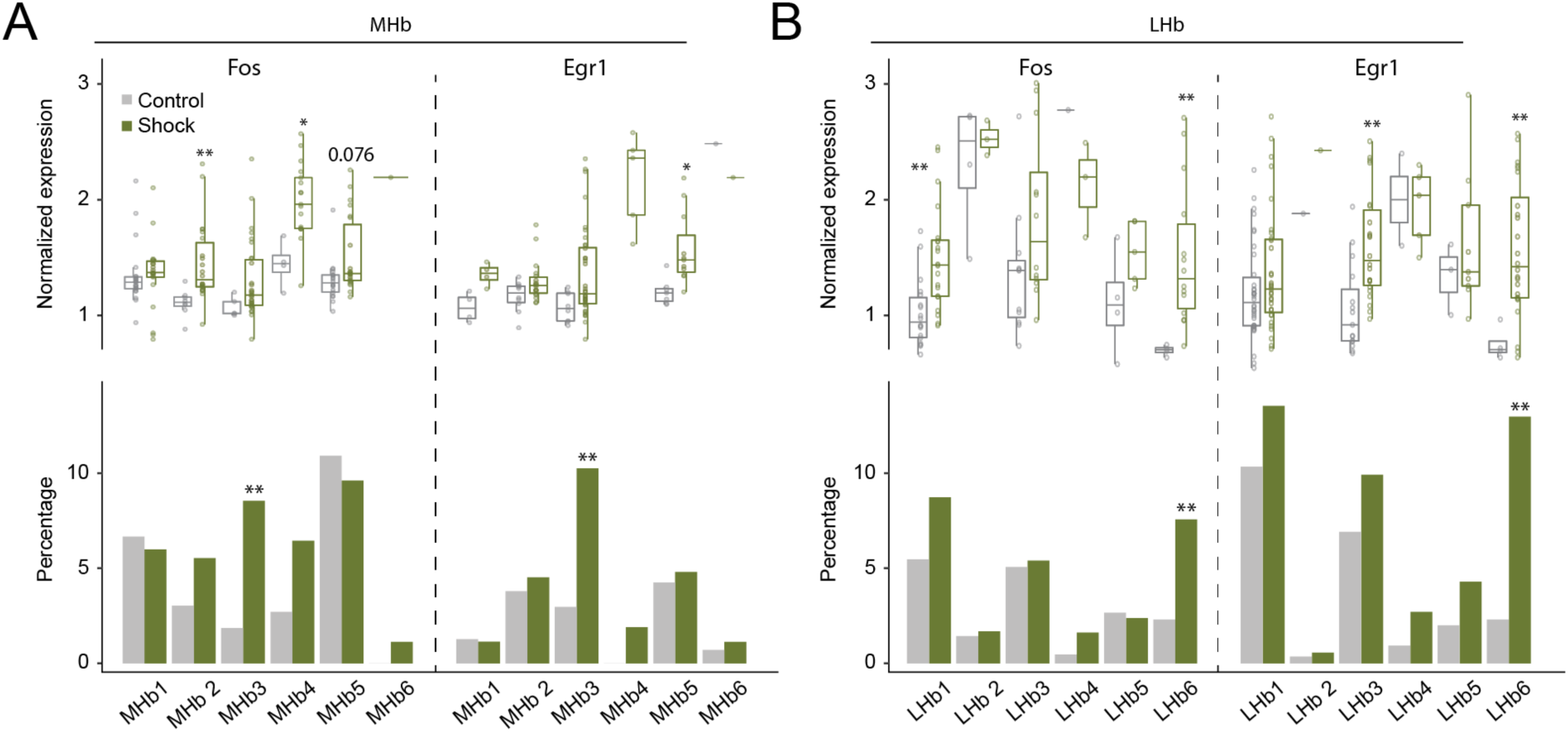
Activation of habenular clusters by aversive stimulus. **A, B.** Comparison of *Fos* and *Egr1* expression values in the expressing cells (Top) and the proportions of expressing cells (Bottom) between control and shock groups in the scRNAseq experiments (**A**: MHb and **B**: LHb). Wilcoxon rank sum test (Top), and Fisher’s exact test (Bottom). *p <0.05, * *p <0.01.

## Discussion

Here we identified transcriptional cell types in the mammalian habenula using scRNAseq and inferred distinct gene regulatory networks in the MHb and LHb. Cross species analysis indicated transcriptional homology between the mammalian MHb and zebrafish dorsal Hb and between the mammalian LHb and fish ventral Hb as previously suggested (Amo et al., 2010). Moreover, Hiplex FISH generally showed consistency with the scRNAseq data in terms of cluster identity, while providing spatial information on the distribution of transcriptionally defined cell types. An aversive stimulus enhanced IEG expression within transcriptionally defined clusters in both the MHb and LHb. These datasets provide important foundational information needed for habenula cell type specific targeting, and for future work where cell type specific gene expression can be leveraged for functional and therapeutic targeting to impact motivational behavioral states.

### Cellular diversity in the mammalian habenula

The habenula comprises anatomically segregated nuclei with distinct connectivity, transcriptional profiles and functionalities (Aizawa et al., 2012; Hikosaka, 2010; Namboodiri et al., 2016). Previous work utilizing immunohistochemistry, *in situ* hybridization and bulk RNA surveying techniques (e.g. microarray) (Amo et al., 2010; Hawrylycz et al., 2012; Le Foll and French, 2018; Wagner et al., 2016) have identified characteristic genes enriched in the habenula of mammals and fish. More recent work by Pandey et al comprehensively and unbiasedly characterized the whole transcriptome of habenula neurons in zebrafish (Pandey et al., 2018). Here we conducted scRNAseq on 11,878 cells including 5,558 neuronal cells in the mammalian habenula and transcriptionally identified 12 neuronal clusters across the MHb and LHb. In the 6 MHb clusters, all of which were marked by *Tac2* expression, *Chat* and *Slc18a3*, classically identified markers utilized for circuit studies targeting the MHb (Zhang et al., 2016), were only expressed in 3 clusters while *Tac1* and *Calb1* were expressed in the orthogonal MHb clusters as previously reported (Yamaguchi et al., 2013). Consistent with reciprocal connections between the LHb and raphe serotonergic system (Amo et al., 2014; Hikosaka et al., 2008; Vertes et al., 1999; Xie et al., 2016), *Htr2c* was enriched in half of the 6 LHb clusters (all were marked by *Pcdh10*), whereas *Slc6a1* and *Cacna2d1* marked the rest of clusters.

Importantly, while all 12 neuronal clusters contained enriched DEGs, few clusters had exclusively selective genes. Rather, the majority of marker genes were discriminative and expressed in multiple clusters as observed in other brain structures using similar approaches (Moffitt et al., 2018; Tasic et al., 2018). This observation motivated us to perform Hiplex FISH to expand conventional 3 color FISH into sequential detections of over 10 discriminative genes covering clusters in the MHb and LHb. Based on the marker genes for clusters identified in scRNAseq analysis, we performed Hiplex FISH to visualize 12 marker genes each for the LHb or MHb and to reconstruct clusters *in situ*. There was largely correspondence between scRNAseq and Hiplex clusters in both the MHb and the LHb (Figure 5). Each scRNAseq cluster was correlated with 1-2 Hiplex clusters. Moreover, Hiplex clusters showed spatially biased distributions (Figure 5). Andres et al anatomically divided the MHb and LHb into subnuclei, based on Nissl staining in rat tissue (Andres et al., 1999). 4 major subdivisions of the MHb were identified comprising superior (MHbS), central (MHbC), lateral (MHbL) and inferior (MHbI) parts while the LHb could be divided into superior (LHbS), parvocellular (LHbMPc), central (LHbMC), magnocellular (LHbMMg), lateral (LHbLO and LHbLMc) and anterior (LHbMA) parts. Our Hiplex FISH analysis suggests that some transcriptionally defined clusters have correspondence with anatomically defined subnuclei while other clusters distribute as part of a subnucleus or across subnuclei. For example, cells in hMHb2 (MHb1), which were enriched with *Spon1,* were in the MHbL, while hMHb4 (MHb3, *Neurod2*) and hMHb5/6 (MHb5, *Cck*) distribute in the ventral part and dorsal/dorsolateral part of the MHbC, respectively. In the LHb, hLHb5 (LHb1) is concentrated in the anterior part of LHbMc while cells in hLHb3 (LHb6, *Sv2c* and *Fam101b*) were located in the ventral part of the LHbLMc and the LHbLO.

### Activation of habenula cell types by aversive stimuli

Previous work has demonstrated that both the MHb and LHb are engaged by a variety of aversive stimuli. To examine this further, we sought to determine whether transcriptionally distinct habenula cell types were preferentially activated (as indexed by IEG expression) following exposure to foot-shock. At least two neuronal cell types (MHb3 and LHb6) displayed an increase in IEG expression following this procedure. It is important to note that we cannot rule out the activation of other habenula cell types following aversive stimulus exposure, as it is likely that other cell types are also activated but perhaps to a lesser degree, or by displaying additional transcriptional differences that were not examined here. Nonetheless, these data point to at least two cell types in the habenula that merit further attention in future studies. In future studies, MHb3 could be genetically targeted using an intersectional approach based on the expression of *Neurod2* and *Lynx1* while LHb6 may be accessible based on the expression of *Sv2c* and *Fam101b*.

These data also suggest that there may be a disconnect between the transcriptional definition of cell types, and their functional encoding dynamics *in vivo* or *ex vivo* following aversive stimulus exposure. Electrophysiological (Amo et al., 2014; Cui et al., 2018; Lecca et al., 2017; Li et al., 2011; Yang et al., 2018) and calcium imaging (Shabel et al., 2019) studies show that large proportions of LHb neurons display an increase in their firing/activity rate in response to, or following aversive stimuli exposure, while our transcriptional profiling results suggests more molecular heterogeneity exists. Thus, it seems likely that despite the striking transcriptional differences across habenula cells, their activity dynamics could be similar (at least in response to simple aversive stimuli exposure), and driven largely by their anatomical connectivity. Future studies exploring how *in vivo* activity dynamics of habenula cell types relates to their molecular features are needed to better resolve this.

### Transcriptional controls on habenular cellular properties

The comparisons of transcriptomes between the whole units of the MHb and the LHb identified 170-289 genes including neurotransmitter related genes, glutamate and GABA receptor related genes, ion channel related genes and morphology related genes, which define cellular properties and functions (Figure 3 and Figure S3). These distinct sets of genes are likely under the unique transcriptional controls in the MHb and the LHb. SCENIC analysis revealed that TFs, which regulates >80% of the DEGs, are distinct between the two nuclei. Moreover, the majority of the DEGs are regulated by the combination of multiple TFs, and some TFs regulate the expression of other TFs in the DEG (e.g. *Tcf7l2* regulates *Nfib* in the MHb) (Figure S3). These results suggest that cellular and functional properties of the MHb and the LHb are maintained by distinct transcriptional networks.

In addition to cellular properties in the adult habenula, the unique patterns of transcription factor activities might regulate the development of patterning and connectivity of the MHb and LHb (Quina et al., 2009). Recently, Lipiec et al found that transcription factor *Tcf7l2* regulates the development of anatomical segregations and axonal patterns of the habenula through *Tcf7l2* knockout assays (Lipiec et al., 2019). Consistent with their findings, our SCENIC analysis showed that *Tcf7l2* regulates a variety of DEGs including ion channels, neurotransmission and cytoskeleton related genes in the MHb and the LHb (Figure S3). The roles of other TFs revealed in our analysis in the habenula development remains elusive and are one of the critical future directions from our study.

### Conservation of cell types across species

The habenula is an ancient and evolutionally conserved structure existing in all vertebrates from fish to humans (Aizawa et al., 2011). The shared features in cytoarchitecture, projection patterns and functions between species have been extensively studied. Raphe projecting neurons in zebrafish are exclusively present in the ventral habenula, which are marked by *Pcdh10a* expression (Amo et al., 2010) and involved in the expectation of danger (Amo et al., 2014), whereas the LHb in mice also projects to the dorsal raphe nucleus (Quina et al., 2015), is highly enriched in *Pcdh10* (Amo et al., 2010) and regulates aversion and reward phenotypes. In contrast, IPN projecting habenula neurons are concentrated in the dorsal part in zebrafish (Aizawa et al., 2005; Gamse et al., 2005) and in the medial part in mice (Herkenham and Nauta, 1979), which are enriched in *Tac2* expression and regulates fear and anxiety related behaviors in both species (Aizawa et al., 2012; Duboué et al., 2017; Ogawa et al., 2012; Yamaguchi et al., 2013; Zhang et al., 2016). Consistent with the previous observations, whole transcriptome analysis of mouse and zebrafish habenula showed that dorsal and ventral parts of zebrafish habenula respectively correspond to the MHb and the LHb in mice. Thus, the habenula is an anatomically, functionally and transcriptionally conserved structure in vertebrates.

Interestingly and consistent with the observations in cross species scRNAseq analysis (Butler et al., 2018; Hodge et al., 2019), clustering without Seurat integration generated nearly non-overlapping mice and zebrafish clusters (Figure 4B and Figure S4), which is likely due to experimental differences between studies (e.g. cellular sequencing depth) as well as non-homologous transcriptomes between species. Structural asymmetry present in zebrafish habenula may partially underlie transcriptional diversity between species (Beretta et al., 2012; Pandey et al., 2018).

### Summary and future directions

While single gene markers and cre driver lines have been employed to gain genetic access to habenula cells (see (Proulx et al., 2014) for discussion), in most cases, the expression of single genes does not accurately reflect habenula cell type identities. This is important to consider for future studies that will employ genetic targeting methods for manipulating habenula cell types, and presents an overall challenge for the field. The transcriptomic analysis of mammalian habenula using scRNAseq and Hiplex presented here derives a census of transcriptional diversity by unbiased clustering, provides comprehensive transcriptomic resources to the community, and infers functionalities of divergent cell types. In addition, there are largely consistent observations between distinct modalities in our dataset (e.g. scRNAseq and Hiplex; mice and zebrafish) and between our analysis and past scholarship, further suggesting the validity of our analysis. The future investigations on the electrophysiological and functional properties of transcriptionally defined cell types are expected to deepen our understanding of the habenula in the complex motivational states.

## METHOD DETAILS

### Animals

C57BL/6J male mice (Jackson Laboratory) were used for single-cell preparation and Fluorescent in situ hybridization (FISH) experiments (P50-55 at tissue extraction). Mice were group housed except that for the last two days before tissue isolation, they were singly housed. Mice had *ad libitum* access to food and water and were kept under a reverse 12 h light-dark cycle. All experiments were conducted in accordance with the National Institute of Health’s Guide for the Care and Use of Laboratory Animals and were approved by the Institutional Animal Care and Use Committee at the University of North Carolina and the University of Washington before the start of any experiments.

### Single-cell preparation, cDNA library construction for scRNAseq

Shock (n=4) and Control (n=4) groups were handled 5 min/day for 7 days. Mice in the Shock group were habituated to the shock chamber (Med Associates Inc.) 10 min/day for 7 days prior to the behavior experiment. On the shock day, mice in the Shock group received 30 foot-shocks (0.3 mA, 1 s duration, mean inter shock intervals: 116 s) in 60 min. Immediately after the shock session, mice were deeply anesthetized with intraperitoneal injection of 0.2 mL of stock solution containing sodium pentobarbital (39 mg/mL) and phenytoin sodium (5mg/mL) and transcardially perfused with ice-old NMDG-aCSF containing inhibitor cocktails which was used throughout procedures unless noted. NMDG-aCSF: 96 mM NMDG, 2.5 mM KCl, 1.35 mM NaH2PO4, 30 mM NaHCO3, 20 mM HEPES, 25 mM glucose, 2 mM thiourea, 5 mM Na+ ascorbate, 3 mM Na+ pyruvate, 0.6 mM glutathione-ethyl-ester, 2 mM N-acetyl-cysteine, 0.5 mM CaCl2, 10 mM MgSO4; pH 7.35–7.40, 300-305 mOsm, oxygenated with 95% O2 and 5% CO2. Inhibitor cocktails: 500 nM TTX, 10 μM APV, 10 μM DNQX, 5 µM actinomycin, 37.7 µM anysomycin. Mice in the control group were removed directly from the home cage. Brains were extracted and coronal sections containing the habenula were prepared at 300 µm using a vibratome (Leica, VT1200). Slices including the habenula (3-4 slices/animal) were recovered in a chamber for 30 min on ice. Habenula samples were then punched (0.75 mm diameter, EMS) from 4 animals/group (6-8 punches/animal) were pooled and enzymatically digested with 1 mg/mL pronase (Roche) for 45 min at room temperature, followed by mechanical trituration with fire-polished glass capillaries (tip diameter 200-300 µm) and filtered through strainers (pore size 40 µm) to remove cellular aggregates. Dead and dying cells in the cell suspension were then removed using a dead cell removal kit (Miltenyi Biotec). After centrifugation, cell concentration was manually counted using a hemocytometer and final cell concentration was adjusted to 1,000 cells/µL.

cDNA libraries were constructed following manufacture’s instruction (Chronium Single Cell 3’ Reagents Kits V2 User Guide, 10x Genomics). Briefly, ∼17,000 dissociated cells were mixed with reverse transcription mix and loaded into the chip. The mRNAs of single cells were captured by barcoded beads using a Chromium controller. Reverse transcribed cDNAs were then PCR amplified, fragmented, and ligated with adaptors followed by sample index PCR. cDNA libraries were sequenced on an Illumina Nextseq 500 (v2.5) and the alignment of raw sequencing reads to the mouse genome was conducted using the 10x Genomics Cell Ranger pipeline (V2) to obtain cell by gene matrices for subsequent downstream analysis.

### Data Analysis for scRNAseq data

Clustering, differential gene expression analysis and integrative analysis of differential conditions, cross species and cross modalities were performed using the Seurat V3 package (Stuart et al., 2019). SCENIC package was utilized to infer gene regulatory networks by transcription factors (Aibar et al., 2017).

### Data preprocessing and doublet removal

Low abundant genes (expressed in less than 3 cells) and cells of potentially low quality (total UMI<700 or total UMI>15,000, or percentage of mitochondrial genes >20%) were removed from downstream analysis. Suspected doublet cells were computationally removed by utilizing DoubletDecon package (Version 1.02.) (DePasquale et al., 2018) with the default settings.

### Integrative Clustering and differential gene expression analysis

To minimize the effects of experimental variations (batch effects and behavioral conditions) on clustering, we used Seurat V3 analysis package (Stuart et al., 2019), which utilized Canonical correlation analysis (Butler et al., 2018) and mutual nearest neighbor analysis (Haghverdi et al., 2018). Briefly, gene counts were scaled by the cellular sequencing depth (total UMI) with a constant scale factor (10,000) and then natural-log transformed (log1p). 2,000 highly variable genes were selected in each sample based on a variance stabilizing transformation. Anchors between individual data were identified and correction vectors were calculated to generate an integrated expression matrix, which was used for subsequent clustering. Integrated expression matrices were scaled and centered followed by principal component analysis (PCA) for dimensional reduction. PC1 to PC30 were used to construct nearest neighbor graphs in the PCA space followed by Louvain clustering to identify clusters (resolution=0.8). For visualization of clusters, Uniform Manifold Approximation and Projection (UMAP) was generated using the same PC1 to PC30.

To identify markers (differentially expressed genes) for neuronal and non-neuronal cells, first, clusters of each cell types (neuron, astrocyte, oligodendrocyte, microglia, ependymal cells, OPC, and Endothelial cells) were combined. And then, expression value of each gene in given combined clusters were compared against the rest of cells using Wilcoxon rank sum test and p-values were adjusted with the number of genes tested. Gene with log fold change >0.25 and adjusted p-value <0.05 were considered to be significantly enriched. To examine the stability and robustness of clustering results, 10-100 % of cells were randomly sub-sampled and clustered by the identical procedure described above for 10 times at each sub-sampling rate.

Based on the expression of canonical markers for neuronal cells (*Stmn2* and *Thy1*), cells in the neuronal clusters were extracted to be re-clustered using the identical integrative clustering and differentially expressed gene analysis described above. Initial clustering resulted in 16 neuronal clusters, among which 4 clusters were excluded in downstream analysis due to their expressions of peri-habenula markers. Those 4 clusters were marked with one or a couple of following genes: *Pdyn*, *Lypd6b*, *Lypd6*, *S1pr*, *Gbx*, *Ramp3*, *Cox6a2*, *Slitrk6* and *Dgat2*. Gene ontology database at Mouse Genome Informatics was used to generate dot plots for the expression of GPCRs (GO: 0004930; G protein-coupled receptor activity; gene names containing “receptor” were used; Figure S2) and neuropeptide (GO: 0007218; neuropeptide signaling pathway; gene names containing “receptor” were excluded; Figure S2.). Similarly, Dot plots illustrating expression of neurotransmitter related genes (GO: 0042133; neurotransmitter metabolic process, GO: 0006836; neurotransmitter transport, GO: 0001504; neurotransmitter uptake, GO: 0007269; neurotransmitter secretion), Ion channel related genes (GO: 0034702; Ion channel complex), and glutamate and GABA receptor related genes (GO: 0008328; Ionotropic glutamate receptor complex, GO: 0098988; G-protein coupled glutamate receptor activity, GO: 0016917; GABA receptor activity (Gpr156 was excluded) were generated for the MHb or LHb enriched genes (Figure 3).

### Gene Ontology analysis

Enriched ontology terms for differentially expressed genes were identified using Enrichr (Chen et al., 2013; Kuleshov et al., 2016). GO Biological Process 2018, GO Molecular Function 2018, and GO Cellular Component 2018 were referenced to identify ontology terms with the adjusted p-value<0.1 (Figure S1 and S3).

### Gene regulatory network analysis

The SCENIC package (Aibar et al., 2017) was utilized to infer transcriptional regulation on the DEGs in the MHb and LHb by transcriptional factors. SCENIC analysis consisted of two major steps: construction of co-expression network using GENIE3 and identification of direct binding by DNA-motif analysis using RcisTarget. Log-normalized expression matrix of habenular neurons generated using Seurat was used as input data. After running GENIE3, motif data set (mm9-500bp-upstream-7species.mc9nr.feather, mm9-tss-centered-10kb-7species.mc9nr.feather) was used to construct regulons for each transcription factor. To examine selectivity of transcriptional regulation on DEGs in the MHb or LHb, selectivity scores for each transcription factor was calculated by dividing the number of DEGs in the regulon by the number of all genes in the regulon. Cumulative plots illustrating the coverage of DEGs by the transcription factors were generated by using sorted transcription factors based on the selectivity score.

### Cross-species analysis

Transcriptomes of Zebrafish and mouse habenulae were integratively analyzed to examine conservation of habenular transcriptome across species. The zebrafish habenula dataset was obtained from the publicly available repository (Accession number: GSE105115). Since our mouse dataset was generated from adult mice using 10x Genomics platform, we also used zebrafish dataset which was generated from adult subjects with 10x Genomics platform (GSM2818522, GSM2818523) to minimize confounding effects introduced by experimental settings. To integrate zebrafish and mouse data, first, gene names in the zebrafish were converted to mouse genes using biomaRt package (Durinck et al., 2005, 2009) and bioDBnet (biological DataBase network). Gene counts of zebrafish genes that were converted to the identical mouse genes were averaged. 8,951 genes were used for integrative analysis. Zebrafish and mouse datasets were integratively clustered using Seurat V3 as described above. 100 variable genes in each sample were used to generate integrative expression matrix. PC1-PC7 were used for clustering and UMAP visualization. To establish correspondence between cross species integrated clusters and mouse or zebrafish clusters, which were generated independently by the clustering approach described in the section of “Integrative Clustering and differential gene expression analysis” (Figure S4), expression values of all cells in each cluster were averaged for each gene and then pairwise Pearson’s correlation coefficient between all pairs of clusters in each dataset was computed. In the heatmaps to visualize Pearson’s correlation of two expression matrix (Figure 4), negative correlations or correlations which were not statistically significant (p>0.05) were adjusted to 0 (White).

### Multiplex/Hiplex FISH

Mice (n=4-5) were deeply anesthetized with isoflurane, decapitated and brains were rapidly flash frozen on dry ice. Coronal sections were cut at 20 µm on a cryostat (Leica) and stored at −80 °C until use. HiPlex and subsequent Fluorescent multiplex assays (RNAscope®) were performed following manufacture’s instruction (Wang et al., 2012). The HiPlex together with Fluorescent multiplex assay allows multiplex detection for up to 15 targets on a single tissue section. Briefly, sections were fixed in 4% paraformaldehyde, dehydrated with 50%, 70%, 100% ethanol, then treated with protease. All the HiPlex probes were hybridized and amplified together. MHb probes; *Neurod2* (#537171-T1), *Spon1*(#492671-T2), *Zmat4*(#578011-T3), *Kcng4*(#316931-T5), *Fn3krp* (#583881-T6), *Ctxn1*(#467041-T7) *Tac2*(#446391-T8), *Col25a1*(#538511-T9), *Cck*(#402271-T10), *Gpr151*(#317321-T11), *Synpr*(#500961-T12): LHb probes; *Necab1*(#428541-T1), *Rflnb*(524091-T2), *Cartpt*(#432001-T3), *Vsnl1*(#583871-T5), *Lmo3*(#497631-T6), *Brinp3*(#583861-T7), *Slc12a5*(#311901-T8), *Pcdh10*(#477781-T9), *Ptn*(#486381-T10), *Gpr151*(#317321-T11), *Gabra1*(#435351-T12). The detection was achieved iteratively in groups of 2-3 targets. After washing, cell nuclei were counterstained with Dapi and samples were mounted. Imaging was performed as described below. After each round, fluorophores were cleaved and samples moved on to the next round of the fluorophore detection procedures. Fluorescent multiplex (ACD Bio) was performed following the HiPlex. Genes targeted with Fluorescent multiplex were as follows; *Slc18a3*(#448771-C3), *Calb2*(#313641-C3), *Hcr2c*(#401001).

Images were obtained with Zeiss ApoTome2 with 20x objective using Zen (blue edition) software. All images from all rounds of staining were then registered to each other to generate ∼15 plex images using HiPlex image registration software (ACD Bio). Cells were segmented and mean fluorescent intensity and XY coordinates of ROIs were measured with image J. Mean fluorescent intensities of ROIs were background-subtracted and thresholds for each gene were applied.

To cluster cells based on the detected mRNAs’ expression matrix and further establish correspondence between clusters in scRNAseq and Hiplex experiments, Seurat V3 was utilized. Cells which did not express any genes were removed for downstream analysis. Expression data were log normalized and scaled, and PCA was conducted using all the genes. PC1-PC10 were used for graph-based clustering and UMAP visualization. Since 2000 variable genes were used for clustering in scRNAseq analysis in comparison to only 12 genes in Hiplex experiments and we also found that our clustering results of scRNAseq data were stable across various parameter settings (data not shown), we assumed that clustering results in scRNAseq were more reliable than Hiplex experiments. Thus, we adjusted the granularity/resolution of clustering in Hiplex experiments to match the numbers of clusters in Hiplex experiments with those of scRNAseq analysis. In the LHb, 1 of 7 clusters was excluded from analysis as it only consisted less than 2 % of all cells. To establish correspondence between clusters in scRNAseq and Hiplex experiments, expression values of all cells in each cluster were averaged for each gene and then pairwise Pearson’s correlation coefficient between all pairs of clusters in each dataset was computed. In the heatmaps to visualize Pearson’s correlation of two expression matrix (Figure 5), negative correlations were adjusted to 0 (White). Since correlation was computed using relatively small number of genes (12), p values were not considered in the heatmap (Figure 5).

### IEG analysis in scRNAseq experiments

Fisher’s exact test was used to test the significance of the proportion of IEG expressed cells in the shock group against the control group. T-tests were used as a statistical test for IEG expression levels. P-values were bonferroni-corrected in all data.

### Data availability

The NCBI Gene Expression Omnibus accession number for the scRNAseq data reported in this paper is GSE137478.

## Acknowledgements

We thank M. Rossi and Stuber lab members for critical comments on the project and the manuscript. We thank C. Trapnell for helpful discussions. We thank S. Ng-Evans for the assistance with behavioral equpment. We thank Y. Tao and Z. Hu (University of North Carolina) for assistance with 10x genomics library generation. We thank R. Ying and O. Kosyk (University of North Carolina) for colony maintenance. This work is supported by the Brain and Behavior Research Foundation (NARSAD Young Investigator Award) (K.H.), the National Institutes of Health (NS007431 (M.L.B.), DA038168 and DA032750 (G.D.S.), the Foundation of Hope (G.D.S.).

## Author contributions

G.D.S. supervised the project. G.D.S. and Y.H. conceived the project, designed experiments, analyzed and interpreted the data, and wrote the manuscript. Y.H. conducted all experiments. K.H. analyzed and interpreted the data, and wrote the manuscript. M.L.B. analyzed the scRNAseq data. Y.L., J.L.S. and O.R.A analyzed the FISH data.

## Figures and Legends

**Figure S1.**
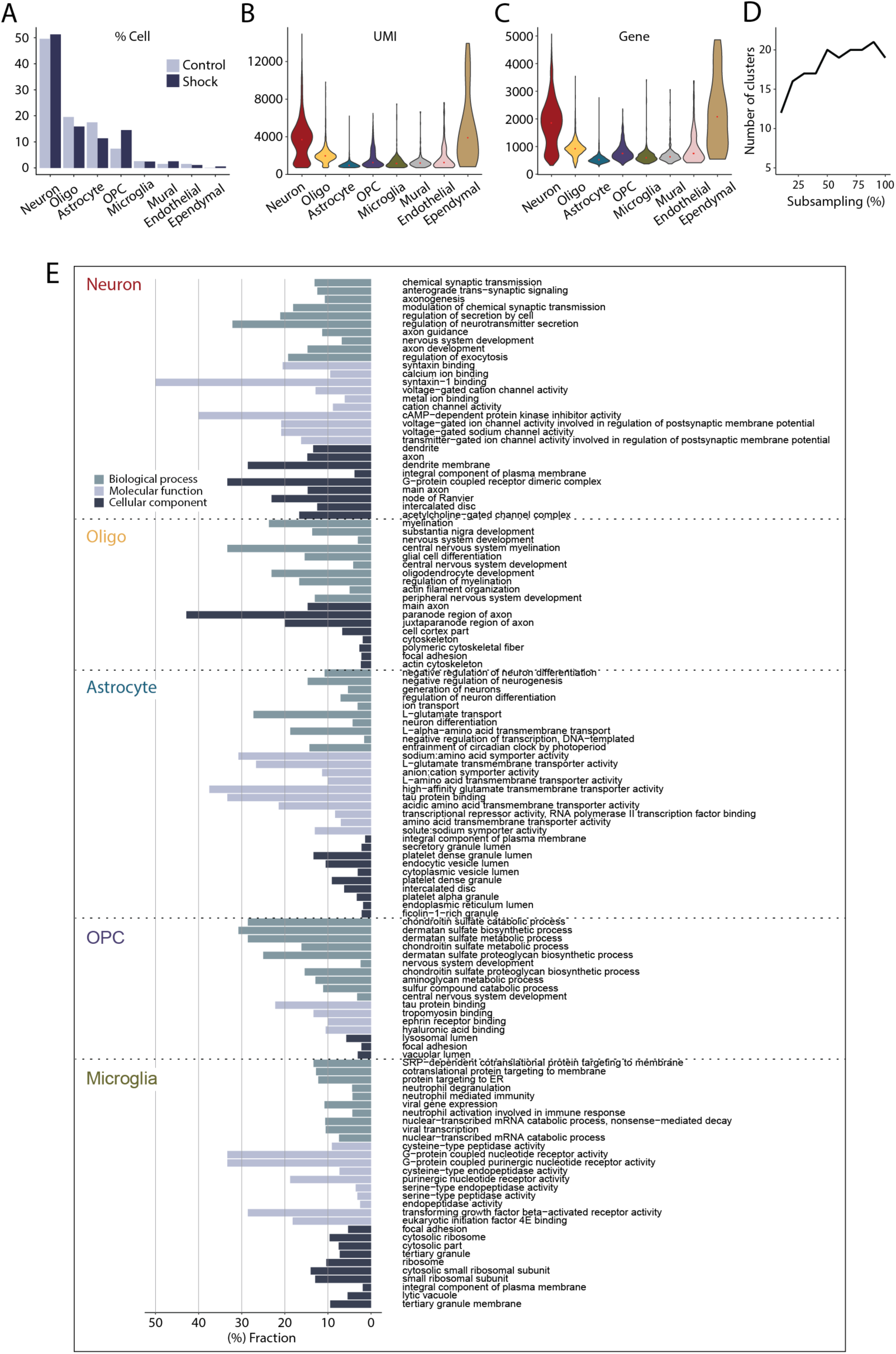
Additional info for scRNAseq experiments, related to Figure 1. **A.** Percentage of cells in each cell type from control or shock groups. **B and C**. Violin plot showing the UMI (**B**) and gene (**C**) distributions in each cell type. **D.** The number of clusters of sub-sampled habenula cells. **E.** Enriched gene ontology terms and fractions of all genes belonging to each term for major cell types. Only terms with adjusted p<0.1 were shown. GO Biological Process 2018, GO Molecular Function 2018, and GO Cellular Component 2018 were referenced.

**Figure S2.**
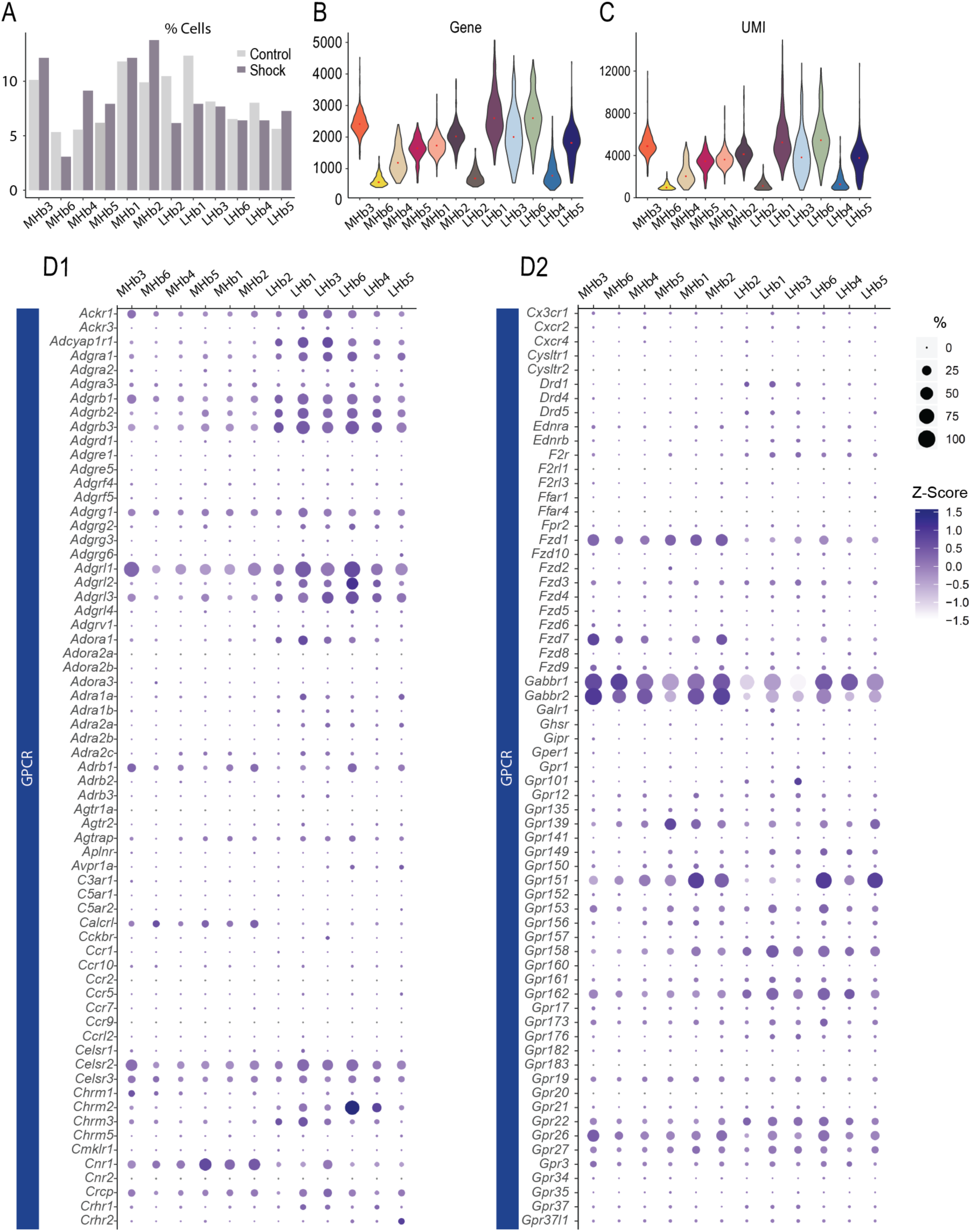
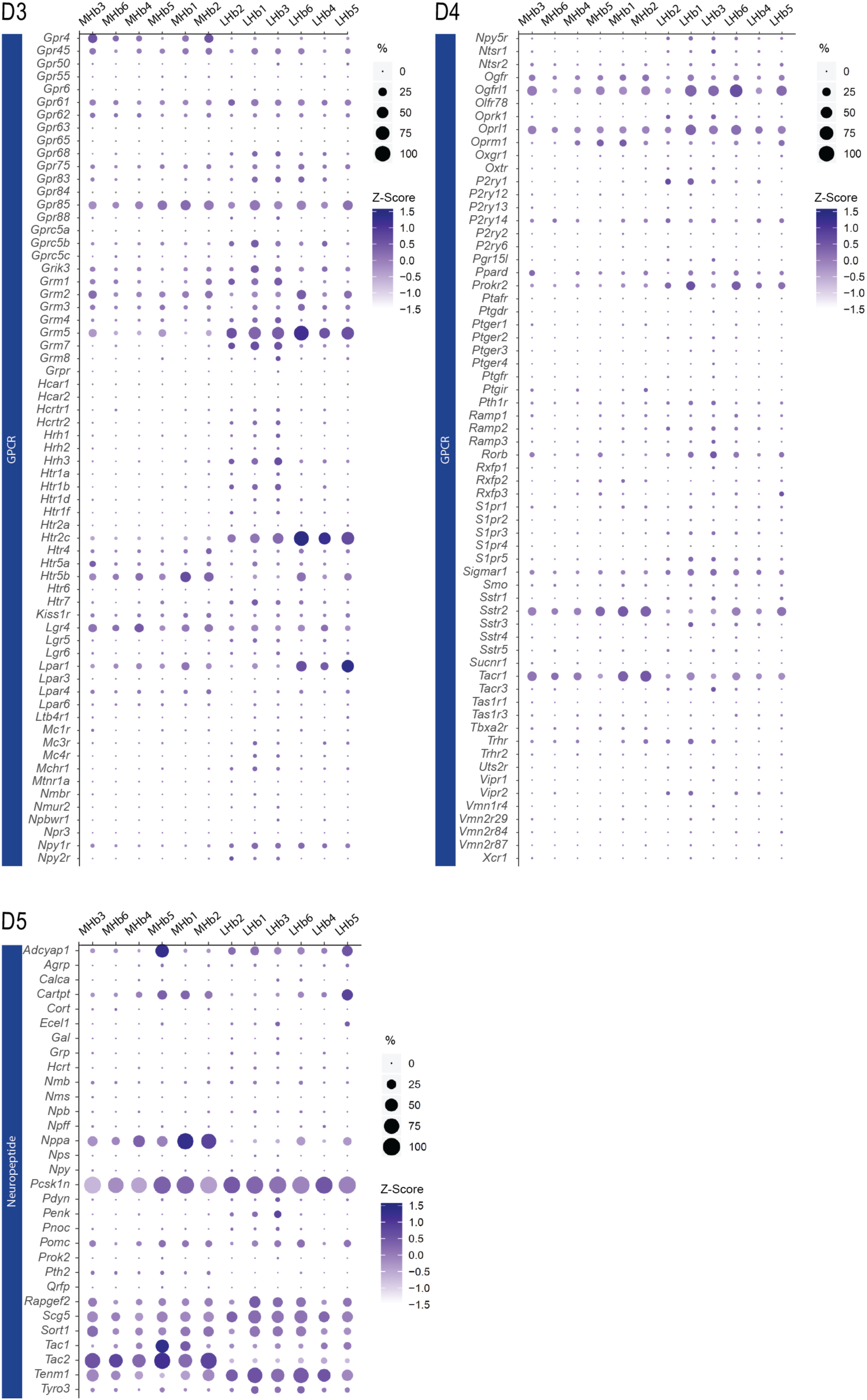
Additional info for scRNAseq experiments on neuronal cells, related to Figure 2. **A.** Percentage of cells in each cell type from control or shock groups. **B and C**. Violin plot showing the gene (**B**) and UMI (**C**) distributions in each cell type. **D.** Dot plot illustrating scaled expression levels (color) and the proportions of expressing cells (dot size) of GPCR and neuropeptide in each cluster.

**Figure S3.**
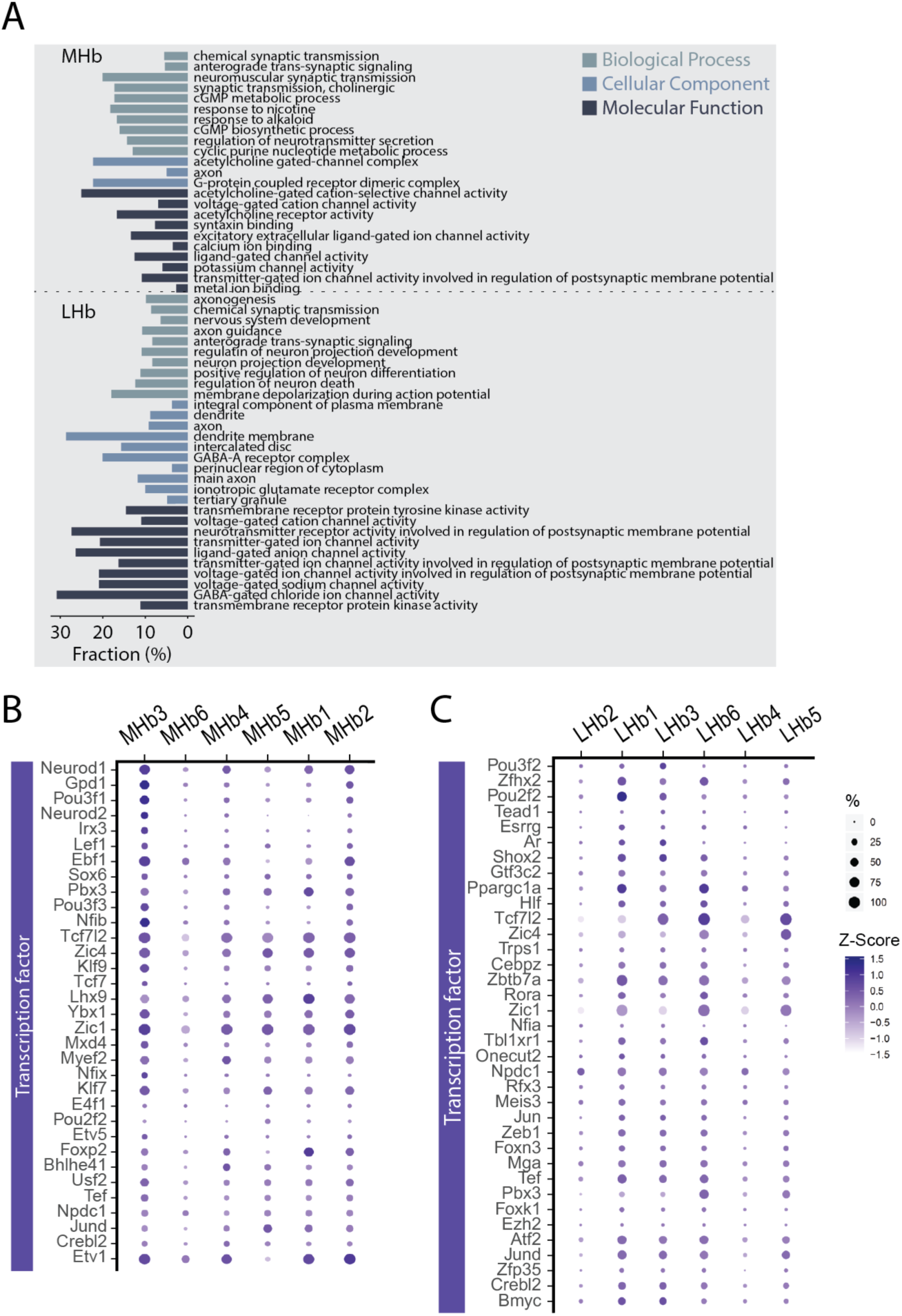
Additional info for scRNAseq experiments on neuronal cells, related to Figure 3. **A.** Enriched gene ontology terms and fractions of all genes belonging to each term in the MHb and LHb. Only terms with adjusted p<0.1 were shown. GO Biological Process 2018, GO Molecular Function 2018, and GO Cellular Component 2018 were referenced. **B and C**. Dot plot illustrating scaled expression levels (color) and the proportions of expressing cells (dot size) of transcription factors in MHb (**B**) and LHb (**C**) clusters. 33 (MHb) and 36 (LHb) TFs, top selective TFs regulating >80% of marker genes, were shown.

**Figure S4.**
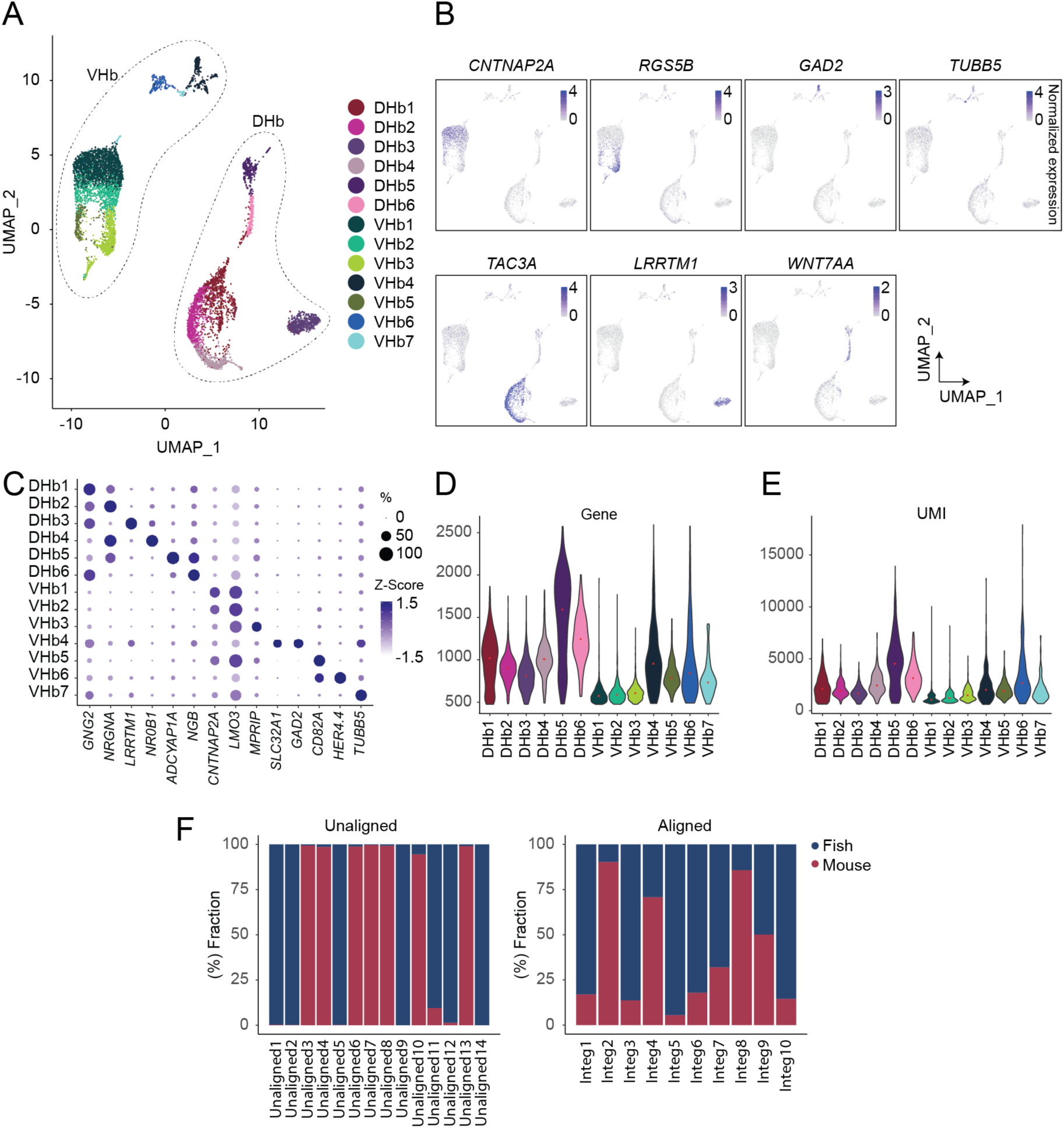
Additional info for scRNAseq analysis on adult zebrafish habenula neurons, related to Figure 4. **A.** UMAP visualization of neuronal clusters of zebrafish habenula. **B.** Expression plots illustrating expression levels of marker genes for dorsal or ventral parts of zebrafish habenula. **C.** Dot plot illustrating scaled expression levels (color) and the proportions of expressing cells (dot size) of discriminative marker genes for clusters. **D and E**. Violin plot showing the gene (**D**) and UMI (**E**) distributions in each cell type. **F.** Proportion of zebrafish or mouse cells in each unaligned (Left) and aligned (Right) cluster.

**Figure S5.**
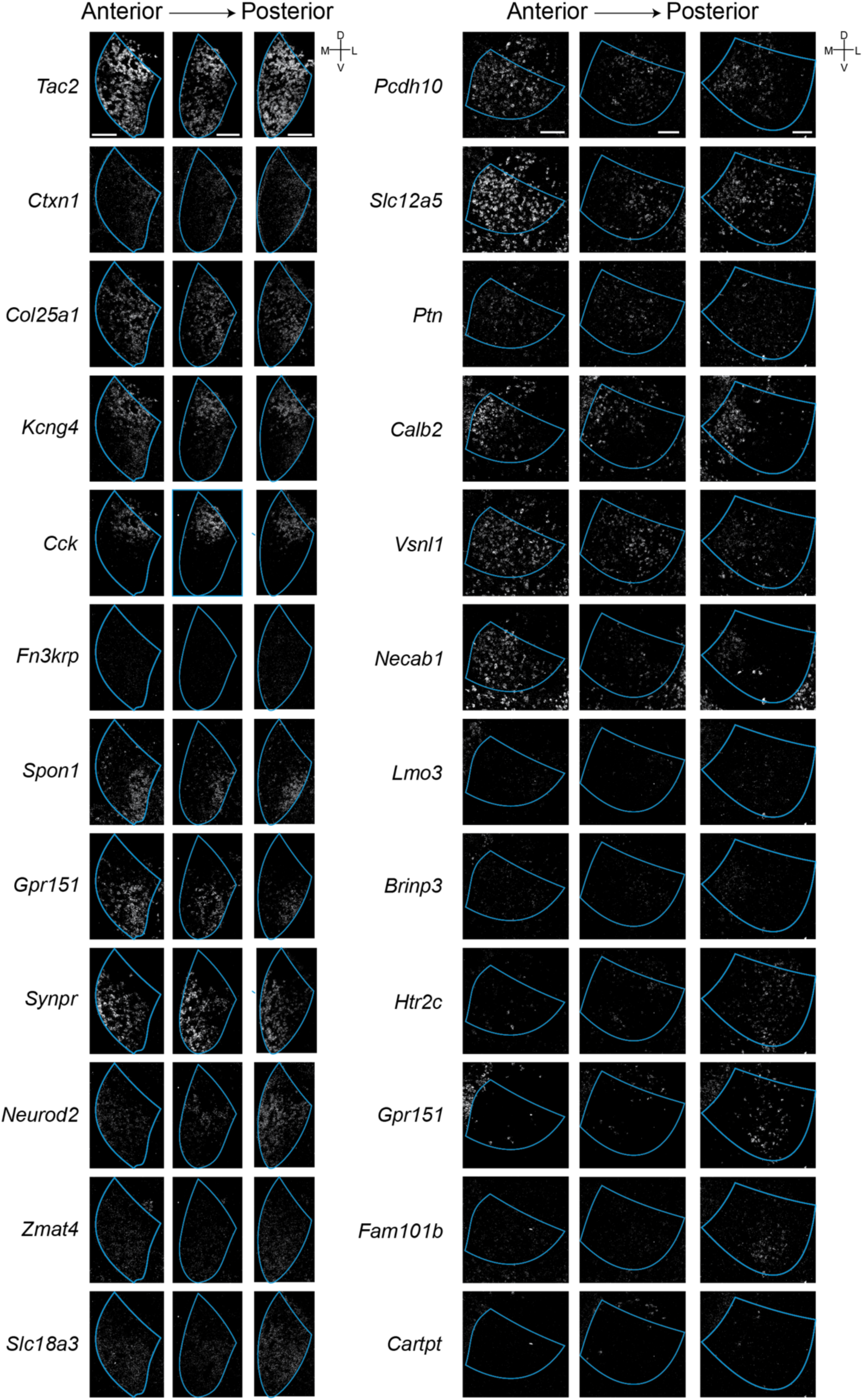
Additional info for Hiplex experiments, related to Figure 5. **A and B.** Representative FISH images of all 12 detected genes for the MHb (Left) and the LHb (Right). Scale bar: 100 µm.

